# Spatio-Temporal Multi-Omics Profiling of Mechanisms and Biomarkers in Inhaled Drug-Induced Lung Toxicity

**DOI:** 10.1101/2025.11.20.689571

**Authors:** Muntasir Mamun Majumder, Eleanor C Williams, Martina Olsson Lindvall, Gregory Hamm, Andrew Jarnuczak, Laura Setyo, Cristina Di Poto, Adnan Azim, Jeff Chen, Anthony A Iannetta, Julia Liz Touza, Erik L Allman, Eric Miele, Julia Lindgren, Elin Sand, Rozita H Anderberg, Steve Oag, Lovisa Franzén, Amanda Costyson, Benjamin P Keith, Jennifer Tan, Stewart Jones, Paul Fitzpatrick, Julia Johansson, Antje Prasse, Dominic Corkill, Annika Borde, Sonja Hess, Sonia Terrillon, Kristoffer Ostridge, Patrik L Ståhl, Per Åberg, Jorrit J Hornberg, Irina Mohorianu, Anna Ollerstam

## Abstract

Comprehensive understanding and early detection of drug-induced lung toxicity remain critical challenges in respiratory drug development. In this study, we propose a multi-omics framework that integrates spatial and temporal tissue-specific transcriptomic signatures with proteomics from minimally invasive biofluids to understand mechanisms and identify safety biomarkers associated with lung toxicity. Using this framework, we identified a panel of candidate biomarkers in bronchoalveolar lavage fluid and plasma, including LCN2/NGAL, RETNLA, SP-D, SPP1/osteopontin, and MMP7, that correlate with histopathological features (e.g., inflammation and epithelial remodeling). We confirmed that these molecular biomarkers were consistently dysregulated across a range of inhaled lung toxicants, human disease (IPF), and environmental exposures (smoke, Alternaria), demonstrating broad applicability across different toxic exposures and translatability. Collectively, this study establishes a robust workflow for mechanism-guided biomarker discovery and proposes a panel of candidates for monitoring drug-induced lung injury in humans.

## Introduction

Inhaled administration of therapeutic drugs is a promising approach to treat respiratory diseases, offering rapid activity and reduced systemic side effects [1, 2]. Nonetheless, drug-induced lung toxicity observed during non-clinical toxicology studies remains a major challenge in the development of novel inhaled therapies; it often manifests as recruitment of immune cells and release of proinflammatory cytokines that may lead to tissue damage [3, 4]. An associated challenge lies in the potential translation and monitoring of these preclinical findings into relevant clinical information. As no predictive safety biomarkers have been identified and validated for this purpose, monitoring for adverse events in the lung traditionally relies on pulmonary function testing, imaging techniques (e.g. computed tomography), or more invasive approaches, including lung biopsies [5]. The former lack sensitivity for detecting effects that precede clinically relevant lesions, thus limiting early intervention [6], while the latter are restricted by their invasiveness, risk of complications, and patient discomfort. To mitigate these limitations, large safety margins are typically applied [7]. As a result, moderate and reversible preclinical outcomes observed at a high dose can restrict the therapeutic dose range in clinical studies, potentially preventing efficacious medicines from reaching patients. Thus, the identification of non-invasive circulating biomarkers predictive of lung injury is critical both for patient monitoring, ensuring patient safety, and for enabling the development of safe and efficacious inhaled therapies[8].

Integrated multi-omics analyses, comprising transcriptomics, proteomics, lipidomics, and metabolomics, especially those with spatial resolution, enable a comprehensive assessment of toxicity mechanisms. These unbiased approaches create opportunities for safety biomarker identification, particularly those mechanistically tied to pathological lesions. The inclusion of spatial omics offers deeper insights into the cellular microenvironment, improves the identification of lung injury-specific gene signatures, and aids in linking these signatures to biofluid markers in bronchoalveolar lavage fluid (BALF) or blood [9]. While several studies have investigated and identified treatment-specific gene expression signatures from the lung, their potential utility as circulatory biomarkers remain unexplored [10, 11].

To address the limited understanding of lung toxicity mechanisms and related safety biomarkers for inhaled compounds, we proposed a spatio-temporal multi-omics analysis framework to profile lung toxicity induced by an inhaled compound with a known toxicity profile in short-term inhalation toxicology studies in rats. Our primary objective was to link histopathological observations with *in situ* drug-induced expression alterations and spatially modulated lipid species, while assessing the detectability of relevant protein biomarkers in biofluids, aiming to identify pathology-associated circulatory biomarkers. We integrated molecular measurements from rat lung tissue with unbiased profiling of BALF to elucidate the underlying mechanisms of lung toxicity and identify functionally linked molecular signature biomarkers in the local lung environment. This mechanistic approach enabled us to uncover key pathways involved in toxicological responses, with some biomarkers subsequently confirmed in plasma, providing insights into systemic manifestations of lung injury mechanisms. To assess the broader applicability of these mechanism-based biomarkers and confirm their ability to reveal common toxicological pathways, we expanded the BALF analysis to include seven additional lung toxicants with diverse mechanisms of action. This cross-model confirmation allowed us to identify shared mechanistic signatures across various contexts of lung injury and assess the mechanistic relevance of the biomarker panel across diverse toxicological scenarios in preclinical species. Mechanistic translational relevance was subsequently assessed and confirmed in human idiopathic pulmonary fibrosis (IPF) lung samples, a disease which shared key pathophysiological mechanisms with drug-induced lung toxicity [12, 13], demonstrating that our approach successfully captures conserved toxicity mechanisms across species and disease contexts.

## Results

### AZ-1 exposure induces dose-dependent lung toxicity

In this study, we utilized AZ-1, a well-characterized small-molecule serine/threonine kinase inhibitor, which had induced lung toxicity of increasing severity in rats at increasing dose levels during the non-clinical development program, to explore toxicity mechanisms and early biomarkers of lung toxicity using tissue and biofluid omics. We conducted a rat inhalation study with two dose levels: 2 mg/kg (lung deposited dose [LDD] 0.2 mg/kg) and 15 mg/kg (LDD 1.5 mg/kg) of AZ-1, and exposure durations of 1, 3, 7 and 14 days (**Fig. 1a**). AZ-1 exposure in the lungs was measured 24 hours after the final dose and confirmed dose- and time-dependent accumulation in tissue, particularly in animals terminated on days 4, 8, and 15 (**Fig. 1b**).

**Figure 1.**
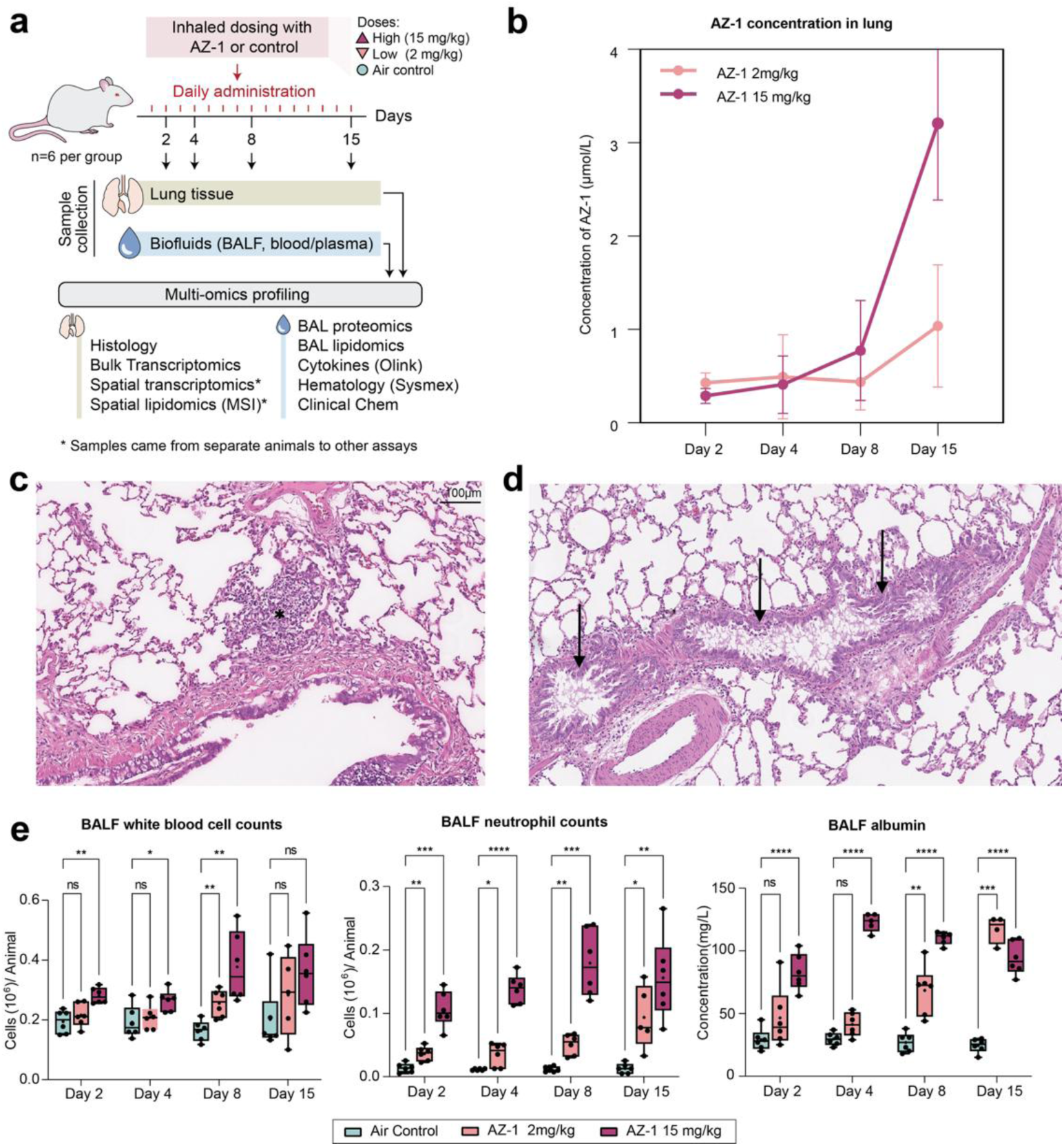
Study overview and investigation of dose- and time-dependent lung toxicity in rats treated with AZ-1. **a.** Overview of the Study design AZ-1 was administered in rats via inhalation at doses of 2 mg/kg/day and 15 mg/kg/day for 14 days. Animals were sacrificed on days 2, 4, 8, and 15 post AZ-1 challenge, with tissue, blood, plasma, and bronchoalveolar lavage fluid (BALF) samples collected for multi-omics characterization, including histopathological analysis, exposure, proteomics, spatial transcriptomics, bulk transcriptomics, and lipidomics. **b.** AZ-1 concentration measured in lung at 24 hours after each treatment. **c**. Multifocal inflammation characterized by the presence of lymphocytes, macrophages, and a smaller number of granulocytes (*) in a representative hematoxylin and eosin image of lung tissue treated with AZ-1 at 15 mg/kg/day and terminated on day 8. **d**. Mild multifocal bronchiolar and alveolar hyperplasia (indicated by arrows) in an AZ-1-treated rat lung at day 8 **e.** Dose- and time-dependent increases in BALF neutrophil counts, white blood cells, and albumin levels in BALF from animals dosed with AZ-1. Significance was assessed using two-way ANOVA followed by Dunnett’s multiple comparison test.

Histopathological evaluation revealed multifocal inflammation, predominantly at the lung periphery (**Fig. 1c, Supplementary fig. 1a**), characterized by neutrophils and macrophages, indicating an active inflammatory response. In affected areas, we also observed type II pneumocyte hypertrophy, a compensatory reaction to lung injury, and disruption of normal lung architecture (**Fig. 1d**). These findings were consistent with the previously described AZ-1 toxicity profile.

BALF analysis supported the histopathological findings, revealing a dose-dependent increase in neutrophils and white blood cells (**Fig. 1e**), along with elevated albumin (**Fig. 1e**) and total protein levels. These changes suggest increased vascular permeability and leakage (pulmonary edema) [14], reflecting disruption of the alveolar-capillary barrier. Together, these findings confirm that AZ-1 induces dose-dependent lung toxicity characterized by inflammation, pneumocyte hypertrophy, and pulmonary edema, providing a foundation for identifying molecular biomarkers linked to these pathologies.

### Spatially differentially expressed genes underline regulatory mechanisms linked to histopathology

To characterize AZ-1-induced lung pathology and identify associated transcriptional signals, we performed spatial transcriptomics (Visium) with H&E-guided annotation of regions of interest (ROI) enriched in cell infiltrates (**Fig. 2a**). Similar ROI were also observed in air control tissues, highlighting the importance of distinguishing treatment-specific expression signatures from background effects.

**Figure 2.**
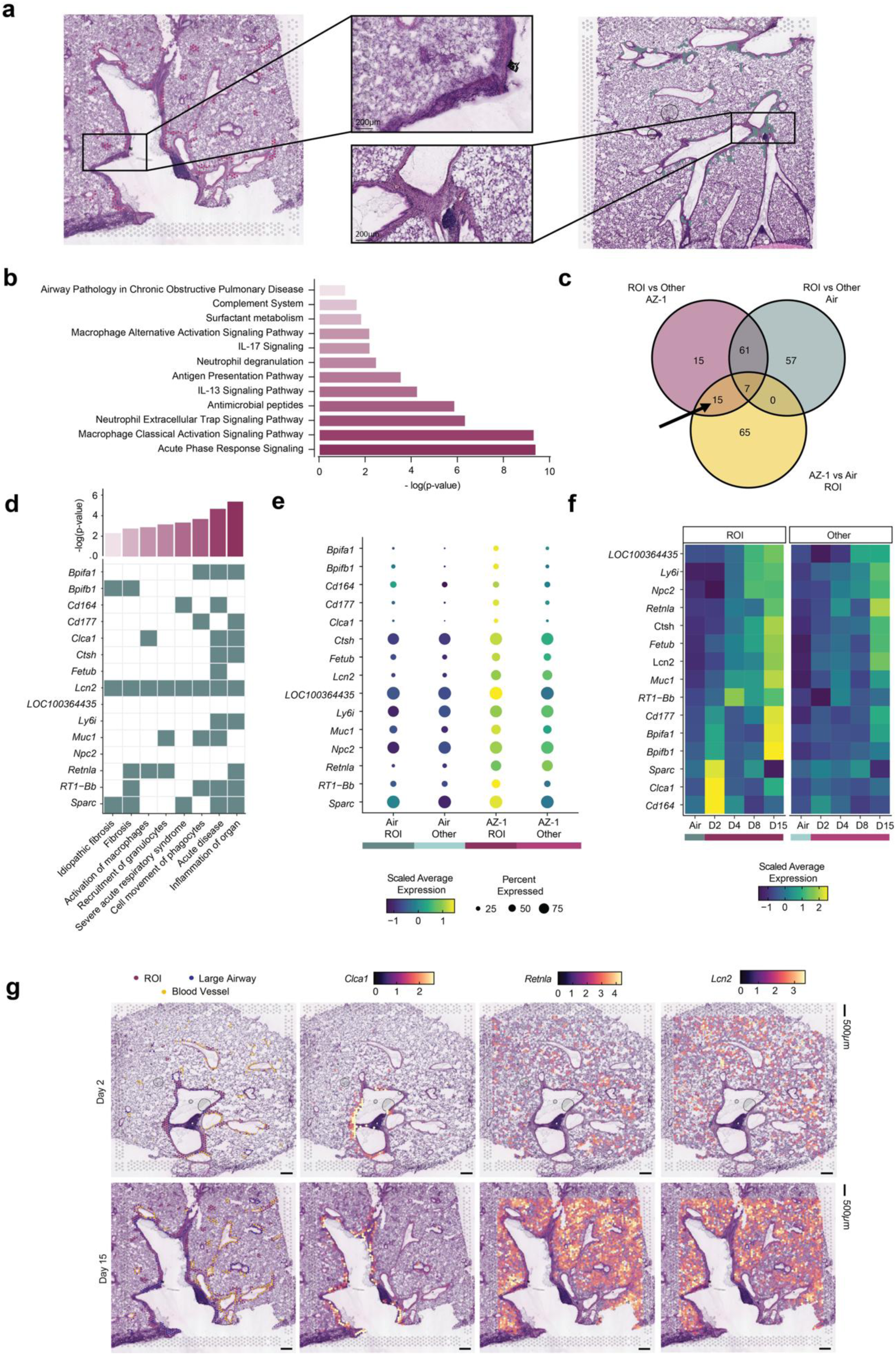
Spatial characterization of differentially expressed genes and associated pathways. **a.** Representative hematoxylin and eosin (H&E) images of lung tissue from an AZ-1 treated (left) and an air control sample (right), annotated to highlight regions of interest (ROI) with histopathological findings, alongside closer views of ROI areas. **b.** Canonical pathways associated with AZ-1 ROI. **c.** Overlap of differentially expressed genes identified in three comparisons: ROI in AZ-1-treated samples vs. ROI in air controls (“AZ-1 vs Air ROI”), ROI in AZ-1-treated samples vs. all other spots in AZ-1 treated samples (“ROI vs Other AZ-1”), and ROI in air controls vs. all other spots in air control (“ROI vs Other Air”). **d**. Canonical pathway associations of the 15 genes of interest. The top bar plot represents significance (−log p-value) for each pathway. **e.** Average expression of the 15 genes differentially expressed exclusively in AZ-1 ROI. Average expression is presented as scaled Z-scores, with dot size indicating the percentage of spots expressing each gene. **f**. Expression patterns (scaled average expression) of the 15 genes of interest across ROI and non-ROI (“other”) regions over treatment duration. **g.** Spatial expression patterns of *Retnla*, *Clca1*, and *Lcn2* in representative AZ-1 treated tissues from two timepoints alongside histopathological annotations. Color scale bars represent normalized expression values

Following initial pre-processing and quality control of the Visium data, differential expression (DE) analysis [15] comparing ROI spots in AZ-1-treated samples *vs.* ROI spots in air controls was performed per timepoint then aggregated to identify treatment-associated changes over time. This comparison revealed distinct transcriptomic expression profiles between the two groups. Pathway enrichment analysis [16] of the differentially expressed genes revealed the canonical pathways impacted by AZ-1 treatment. In alignment with the observed histopathological features, we found significant enrichment of immune and lung-specific processes in AZ-1-treated samples, with the top canonical pathways including acute phase response signalling, macrophage activation, neutrophil extracellular trap signalling and degranulation, IL-13 and IL-17 signalling, and airway pathology in chronic obstructive pulmonary disease (COPD) (**Fig. 2b**).

To further refine the analysis and pinpoint genes specifically linked with ROI in AZ-1 treated samples, we performed two additional DE analyses: (1) ROI in AZ-1-treated samples *vs.* all other regions in AZ-1 treated samples, “AZ-1 other”, and (2) ROI in air control *vs.* all other regions in air control, “Air other”. By intersecting the results (significant in at least one timepoint) from these comparisons with the initial ROI-based analysis (ROI in AZ-1-treated vs. ROI in air control), we identified 15 genes specifically linked to AZ-1 treatment-induced histopathological changes (**Fig. 2c-f**). This approach ensured we captured expression changes most strongly associated with lung toxicity, while excluding baseline differences in air control tissues. These 15 genes represent key molecular signatures of AZ-1-induced histopathological changes and subsequently potential biomarker candidates indictive of treatment-induced lung pathology.

Pathway analysis of the identified genes revealed enrichment in key processes related to lung injury and inflammation (**Fig. 2d**). These included acute disease (*Bpifa1, Cd164, Clca1, Ctsh, Fetub, Lcn2, Ly6i, Muc1, RT1-Bb, Sparc*) and inflammation (*Bpifa1, Cd177, Clca1, Ctsh, Lcn2, Ly6i, Retnla, RT1-Bb, Sparc*) pathways, indicating a strong initial immune response in the affected tissue regions. Additionally, enrichment of pathways associated with fibrosis (*Bpifb1, Lcn2, Retnla, RT1-Bb, Sparc*) and macrophage activation (*Clca1, Lcn2, Retnla*) suggests ongoing tissue remodelling and chronic inflammatory processes beyond the acute phase. Taken together, the involvement of both early- and late-phase pathways and the consistently higher expression of these genes in AZ-1 ROI spots compared to other tissue regions support their potential utility for monitoring different stages of lung toxicity and underscore their association with treatment-induced pathological changes (**Fig. 2e**)

Further analysis of temporal expression trends revealed that most of the 15 genes exhibited a time-dependent increase in expression in AZ-1 ROI, with progressively higher expression at later timepoints, making them potential biomarker candidates for lung injury progression. In contrast, a subset of genes (*Sparc*, *Clca1*, and *Cd164*) showed AZ-1 ROI peak expression in the earliest timepoint, suggesting their involvement in early phase toxicity, indicative of initial lung damage and inflammation (**Fig. 2f**). These genes are linked to pathways associated with immune cell recruitment and early tissue remodelling. Specifically, *Sparc* is well-known to promote macrophage M2 polarization and extracellular matrix remodelling [17, 18], *Clca1* supports macrophage activation and chemokine-driven immune infiltration to the respiratory epithelium [19, 20], and *Cd164* regulates chemokine-mediated immune cell activation and trafficking [21–23].

The visualization of spatial expression gradients revealed diverse patterns. While many genes, such as *Clca1* showed high expression in ROI, others, including *Retnla* and *Lcn2*, were more diffusely expressed but still elevated near ROI in treated samples (**Fig. 2g**). Genes like *Ctsh* and *LOC100364435* exhibited broad expression across the tissue, indicating roles that may extend beyond lesion-specific responses.

### Spatial gene expression and cell patterns in regions of lung injury

To explore how AZ-1-induced toxicity affects cell composition and corresponding biomarker expression, we performed cell-type deconvolution using a mouse single-cell RNAseq reference dataset [24]. We analysed inferred cell type densities across the four groups (“AZ-1 ROI”, “AZ-1 other”, “Air ROI”, and “Air other”) (**Fig. 3a**). The analysis revealed distinct patterns of marker gene expression between conditions, reflecting shifts in cell populations linked to treatment and injury. Interstitial macrophages were most abundant in AZ-1 ROI, indicative of recruitment to sites of injury. Similarly, type 2 pneumocytes showed highest abundance in AZ-1 ROI, suggesting active epithelial repair at the site, whereas type 1 pneumocytes were enriched in the “Air other” group, reflecting normal alveolar structures.

**Figure 3.**
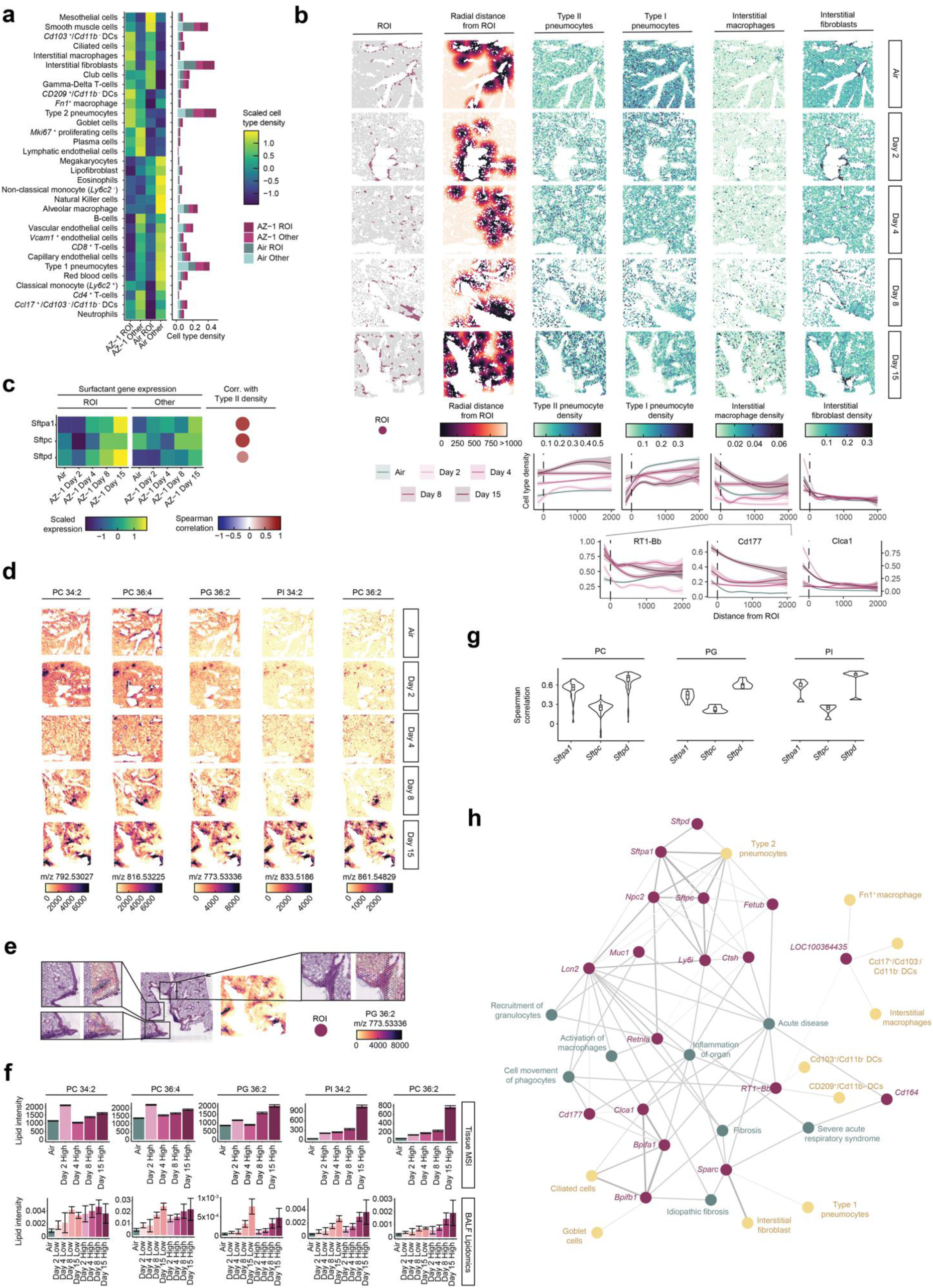
Spatial transcriptomics, cell type dynamics, and lipidomics in AZ-1-treated lungs. **a.** Variations in inferred cell type density across treatment and annotation groups. Color scale of the heatmap represents scaled cell type density scores and the bar plot shows the unscaled cell type density scores in each treatment and annotation group. **b**. Spatial visualization of inferred cell type densities of type II and type I pneumocytes, interstitial macrophages, and interstitial fibroblasts (capped at 99th percentile and with transparency of points scaling with density), shown in relation to ROI annotations and radial distance from ROI. Line plots display cell type scores and scaled expression of selected genes (*RT1-Bb*, *Cd177*, and *Clca1*) as a function of radial distance (0-2000 um) from the ROI, separated by timepoint. Dotted vertical lines mark the ROI border. **c**. Scaled expression of *S*ftpa1, *Sftpc*, and *Sftpd* in ROI and non-ROI (“Other”) regions across conditions. On the right-hand side, the Spearman correlation between each surfactant gene and inferred type II pneumocyte density scores is shown, where dot size and color represent correlation strength. **d**. Spatial distribution of selected lipid species (PC 34:2, PC 36:4, PG 36:2, PI 34:2, and PC 36:2) across representative samples from each condition. Color scale represents the relative ion abundance (m/z peak intensity) in the MSI data. MSI images were co-registered with the spatial transcriptomics datasets, and the same samples are shown as in panel b. **e.** Spatial view highlighting areas of overlap between MSI hotspots in PG 36:2 and ROI annotations after co-registering MSI and spatial transcriptomics data. **f.** Bar plots (with 95% confidence bars) showing changes in intensity for selected PC, PG and PI lipid species found in common between BALF LC-MS lipidomics and tissue mass spectrometry imaging (MSI). **g.** Variation in Spearman correlation between surfactant protein expression in bulk tissue transcriptomics data with PC, PG and PI species which showed significant differences between air and 15mg/kg AZ-1 samples in BALF lipidomics data from the same animals. Violin and box plots capture the variation in correlation across lipids within the same class. **h.** Gene regulatory network showing connections between the 15 ROI-specific genes as well as surfactant genes, their associated pathways, and inferred cell types. Pink nodes represent the 15 AZ-1 ROI-specific DEGs and surfactants, yellow nodes represent inferred cell types, and green nodes highlight enriched canonical pathways. Edges between genes and cell types correspond to the top weighted edges using GENIE3, while edges to pathways were introduced where a gene was annotated as contributing to that pathway.

To assess spatial shifts in lung cell composition, we analysed the inferred cell-type density across a radial distance from the annotated ROI per timepoint and condition (**Fig. 3b**). Interstitial fibroblasts and macrophages exhibited the highest densities at ROI, decreasing with distance. Fibroblast abundance peaked at day 2, consistent with early tissue remodelling, while interstitial macrophages were most enriched at day 15, indicating a more chronic inflammatory response. Interestingly, *Clca1*, a gene involved in macrophage activation and airway inflammation [25], followed a pattern aligned with fibroblast dynamics, with peak expression at AZ-1 ROI on day 2. This suggests that *Clca1* may contribute to the early inflammatory response, potentially priming macrophage recruitment and activation in later stages of injury. Similarly, *Cd177* and *RT1-Bb*, both associated with immune activation, showed high expression at AZ-1 ROI, decreasing with distance. This pattern mirrored the enrichment of interstitial macrophages, particularly at later timepoints, supporting their potential as immune-related biomarker candidates.

Type 2 pneumocyte density increased over the treatment course in AZ-1-exposed lungs, whereas type 1 pneumocytes were depleted in the ROI but maintained in more distal regions. Unlike type 2 cells, type 1 pneumocytes were overall more prevalent in air control samples, suggesting that AZ-1 disrupts normal alveolar epithelial integrity, leading to type 1 depletion and compensatory type 2 expansion.

Given this shift, we next examined the expression of *S*ftpa1, *Sftpc*, and *Sftpd*, which encode surfactant proteins primarily expressed by type 2 pneumocytes, across conditions (**Fig. 3c**). All three genes were upregulated in ROI with AZ-1 treatment, particularly at later timepoints. *S*ftpa1 was upregulated at days 4, 8 and peaking at day 15. *Sftpc* had highest expression at days 8 and 15, with minimal expression at day 2. Similarly, *Sftpd* showed enrichment in AZ-1 ROI, peaking at day 15. In non-ROI (“Other”) areas, the overall trend was similar, although less pronounced. These expression patterns correlated strongly with inferred type 2 pneumocyte density (**Fig. 3c**). While these genes were not identified in prior DE analyses, the increase in expression in AZ-1 samples and in ROI and their established association and correlation with type 2 pneumocytes suggests that their upregulation corresponds to the observed type 2 pneumocyte expansion in response to treatment, supporting their potential as biomarker candidates.

### Surfactant lipid changes reflect spatial lung pathology

To further explore epithelial responses and the role of surfactant metabolism in AZ-1-induced lung toxicity, we next integrated spatial transcriptomics with spatial lipidomics using mass spectrometry imaging (MSI). Spatial lipidomics analyses revealed local accumulation of key surfactant-associated phospholipids including phosphatidylcholine (PC), phosphatidylglycerol (PG) and phosphatidylinositol (PI) (**Fig. 3d**). At days 8 and 15, these lipids showed a distinct clustered distribution aligned with histopathological findings, in contrast to a more diffuse pattern at day 2 and day 4 (**Fig. 3e**). This temporal trend paralleled the upregulation of surfactant genes in type 2 pneumocytes, suggesting a transcriptional-lipidomic coupled response to AZ-1.

To assess whether tissue lipid changes extend into biofluids, we analyzed BALF from AZ-1-treated animals. The spatial lipidomics data (at pseudo-bulk level) showed correlation with dose- and time-dependent surfactant lipid increases in BALF (**Fig. 3f**). In the high-dose samples, alignment between MSI and BALF lipidomics was especially pronounced across all phospholipid species, particularly after 7 days of AZ-1 treatment (7 surfactant-associated lipids significantly modulated in BALF were also detected by tissue MSI). We further assessed the relationship between the main phospholipid classes in BALF (PC, PG and PI) and surfactant tissue gene expression from the same animals (**Fig. 3g**). Significant correlations were observed, with *Sftpd* exhibiting the highest correlation with key pulmonary surfactant phospholipids, followed by *S*ftpa1 and *Sftpc*.

Together, these results demonstrate that surfactant-associated gene expression and lipid species co-localize within spatially defined lung lesions (**Fig. 3c**). These molecular signals, detectable in BALF, support using surfactant-related lipids and proteins as biofluid biomarkers of drug-induced lung toxicity.

To further contextualize the coordinated changes observed in regions of injury, we constructed a gene regulatory network (GRN) [26] connecting key genes to their associated pathways, and inferred cell type correlations (**Fig. 3h**). Genes such as *Lcn2*, *Retnla*, *Ly6i, Npc2*, *Ctsh*, *Fetub, Sftpd, S*ftpa1*, and Sftpc* emerged as key nodes/regulatory hubs with multiple associations with inflammatory processes, epithelial-associated activity, and fibrosis, suggesting a central role in AZ-1-induced pathology. Several genes were linked to macrophage activation, granulocyte recruitment, and epithelial remodelling, highlighting their roles in both acute and chronic phases of lung toxicity. *Lcn2* and *Retnla* were connected to fibrosis, macrophage activation, and type 2 pneumocyte accumulation, indicating their involvement in both inflammation and tissue remodelling. Similarly, *Clca1* and *Cd177* were associated with immune infiltration and phagocyte recruitment, aligning with their spatial enrichment in regions with histopathological changes.

Of the total 18 spatially enriched genes identified, 15 are predicted to encode secreted proteins detectable in circulation based on the Secreted Protein Database (SEPDB) [27], making them potential candidates for circulatory biomarkers. Altogether, these findings suggest that AZ-1-induced lung toxicity is characterized by an interplay between immune activation and epithelial remodeling, and lead to the identification of 15 spatially resolved markers expressed at the site of histopathological lesions and detectable in biofluids.

### Multi-omics analyses reveal mechanistic biomarkers of AZ-1-induced toxicity and underlying pathways

Histopathological evaluation of spatial transcriptomics sections revealed regions with cell infiltrates but was not able to detect diffuse edema or multifocal type 2 pneumocyte hyperplasia, which are subtle and often not detectable in frozen tissue sections. To overcome this limitation and more comprehensively capture such alterations, we conducted transcriptome-wide analysis of whole lung tissue. This allowed detection of gene expression changes associated with edema and hyperplasia that were not visible in the spatial transcriptomics samples. By comparing control and AZ-1 treated lung tissues at both low and high doses across four time points, we uncovered differentially expressed genes and pathways relevant to AZ-1-induced lung toxicity.

AZ-1 treatment markedly modulated the lung transcriptome, particularly in the high-dose group, where 65 genes were significantly differentially expressed, including 58 upregulated genes (adjusted p < 0.05 and log2(FC) > 1.5). In the low-dose group, eight genes were differentially expressed, including four upregulated genes (*Lcn2*, *Retnla*, *Cd177* and *Chia*) (**Fig. 4a**). Functional annotation of the upregulated genes revealed that 51% were associated with cellular or leukocyte migration and 23% were involved in inflammatory responses, including *Chi3l1, Clca1, Cxcl5, Lcn2, Retnla,* and *Spp1*, which play roles in phagocyte activation and neutrophil chemotaxis. Cell cycle and proliferation pathways were also enriched in the high-dose group, with similar patterns emerging at later time points in the low-dose group. This proliferation likely underlies the hyperplasia observed with this compound. Together the gene expression signature was linked to, and revealed, factors associated with AZ-1-induced inflammation and hyperplasia.

**Figure 4.**
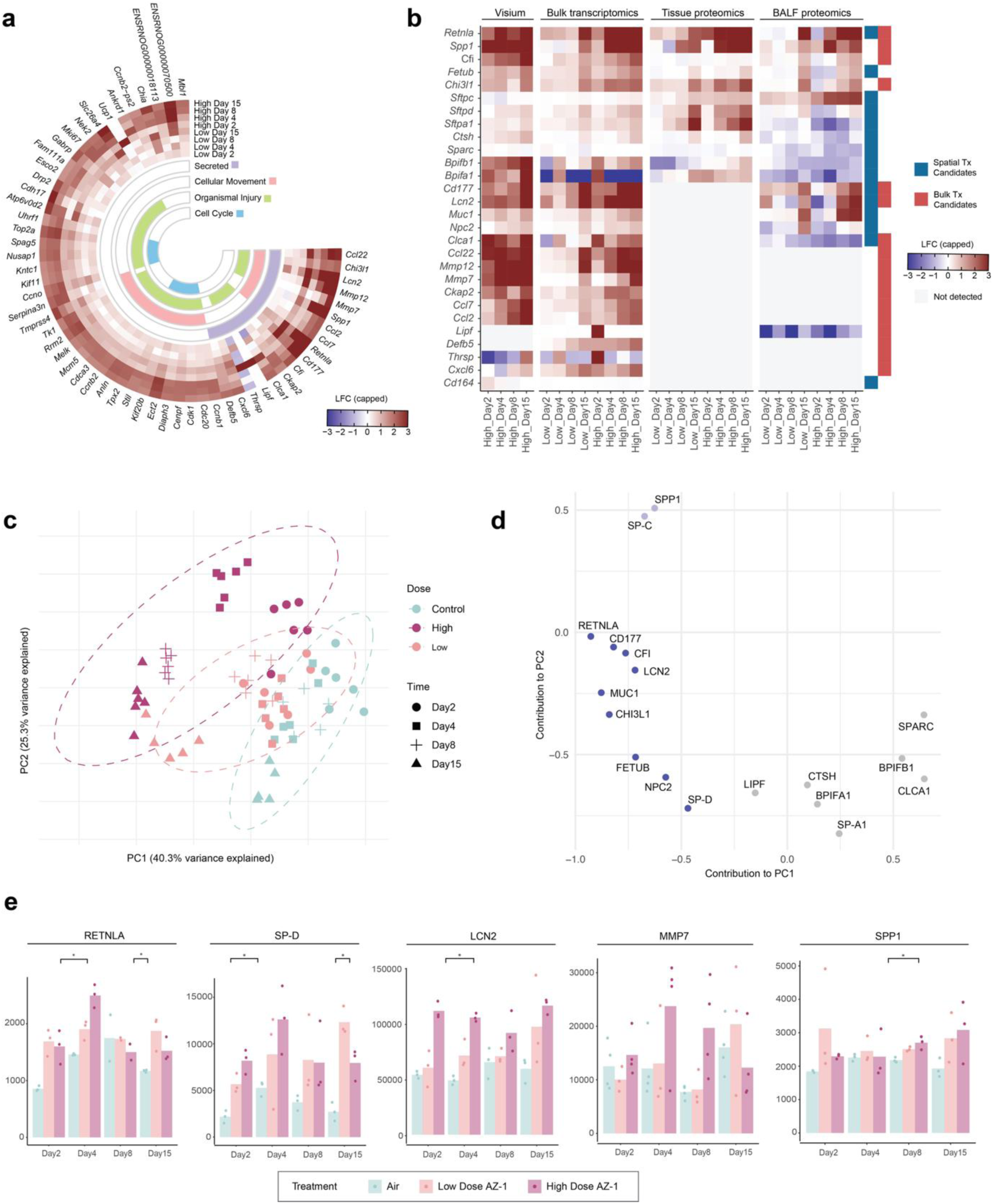
Gene expression and protein detection in AZ-1-treated lung tissue, BALF, and plasma. **a**. Heat map of the most significant gene alterations with AZ-1-dosed groups across timepoints. Heat map displays relative fold-change (log2FC) and shows only hits differentially expressed with log2FC >1.5 between air control and high dose and BH adjusted p-value < 0.05. Selected genes were grouped according to their biological functions. Differentially up- and down-regulated mRNAs are indicated in red and blue, respectively. **b**. Summary heatmap illustrating genes identified in tissue using spatial and bulk transcriptome data, alongside their protein products detected in tissue and secreted in BALF. **c.** Principal component analysis (PCA) dimensionality reduction showing three clusters separating treatment conditions from control across days **d.** PCA loadings plot displaying the contributions of individual proteins to the first two principal components (PC1 and PC2). Proteins selected for further analysis are shown in blue, with potential early biomarkers highlighted in light blue. **e.** Protein levels of RETNLA, SP-D, LCN2, MMP7, and SPP1 in plasma samples from AZ-1-treated and air control animals (Kruskal-Wallis test with Dunn’s post-hoc test using Benjamini-Hochberg multiple-testing correction).

Among the differentially expressed genes, 17 encoded secreted proteins predicted to be detectable in circulation according to SEPDB [27, 28]. These included four candidates, *Lcn2*, *Cd177, Clca1* and *Retnla*, also detected at the histopathology annotated ROI using spatial transcriptomics and 13 additional candidates, *Spp1, Chi3l1, Cfi, Ccl2, Ccl7, Ccl22, Cxcl6, Ckap2, Thrsp, Lipf, Defb5, Mmp7 and Mmp12,* which were included as potential circulatory candidates.

A total of 28 biomarker candidates were curated from spatial and bulk tissue analysis: 15 from spatial gene and lipid analysis (including genes encoding SP-A, SP-C and SP-D) and 13 additional genes from bulk transcriptomics. We then investigated whether these candidates were detectable at the protein level in tissue and BALF, supporting their potential as secreted, circulatory biomarkers. Figure 4b summarizes the overlap of candidate molecules identified across spatial transcriptomics, tissue transcriptomics, and proteomics datasets from tissue and BALF, showing that most spatially identified genes were detected in BALF proteome.

To evaluate predictivity and specificity of these candidates, we assessed the separability of classes on a PCA built on BALF proteomics data using the 18 of the 28 candidates detected in our dataset. Three distinct clusters were observed, effectively separating air control from the high-dose group, with the first principal component (PC1) explaining 40.3% of the total variance (**Fig. 4c**). Among the high-dose samples, we observed two clusters: one encompassing the earlier timepoints (day 2/day 4), and one comprising the later timepoints (day 8/day 15). Interestingly, the late timepoint high-dose cluster overlapped with the day 15 low-dose samples, aligning with histopathological and BALF neutrophil enrichment findings that suggest a toxicity response.

We next examined the principal component loadings to identify the specific proteins contributing to the separation between air control and high-dose animals (**Fig. 4d**). SPP1 and SFTPC exhibited high positive loadings in PC2 and high negative loadings in PC1, aligning with earlier timepoint high-dose samples. Several other proteins showed strong negative loadings in PC1, corresponding to later timepoint high-dose samples. We selected all proteins with a loading < −0.25 in PC1 as potential candidates for separating high-dose samples from controls, which yielded 12 proteins. To confirm this set, we performed targeted partial least-squares discriminant analysis (PLS-DA) which identified a first component separating air control, low-dose, and high-dose groups, and a second component separating timepoints within the high-dose group. We observed a full overlap between the proteins that showed positive correlation with the first component in the PLS-DA analysis and the 11 proteins identified from the PCA analysis; SPP1 and SFTPC additionally showed positive correlation with the second PLSDA component which confirms their potential as early biomarker candidates in BALF.

To investigate potential for minimally invasive detection, we evaluated whether these candidate biomarkers could be detected in plasma from AZ-1-treated animals. ELISA kits for detection in rat plasma were available for LCN2, RETNLA, MUC1, SPP1, SP-A and SP-D. Although MMP7 was not initially detected in BALF proteomics, ELISA confirmed its presence in both BALF and plasma, justifying its inclusion as a known lung injury biomarker [29, 30].

In line with the BALF proteome data (**Fig 4b**), RETNLA levels increased in plasma on day 4 in the high-dose group (**Fig. 4e**). On day 2, plasma concentrations of RETNLA also approximately doubled from a baseline of 856 ± 46 pg/ml to 1686 ± 235 pg/ml (low dose) and 1599 ± 295 pg/ml (high dose). SP-D showed significant upregulation in the high-dose (8.2 ± 1 ng/ml) group compared to air control (2 ± 0.6 ng/ml) on day 2 and remained high during the 15-day follow-up period. Similarly, mean LCN2 levels were elevated in the high-dose group with 111 ± 4 ng/ml on day 2, compared to the air control of 54 ± 1.8 ng/ml. A significant increase in the low-dose group for LCN2 was observed on day 4, and on day 15 mean LCN2 level rose to 9.7 ± 2.3 ng/ml vs. 5.9 ± 6.5 ng/ml in controls. Plasma MMP7 levels increased in the high-dose group on days 4 and 8, and in the low-dose group on day 15. BALF MMP7 was also measured, with samples showing a 9-fold change after a single dose and up to a 75-fold change on day 8 for the high-dose group. For the low-dose group, the BALF MMP7 level changed only on day 15 with a 25-fold increase (5 ± 2.2 ng/ml to 13.7 ± 3.7 ng/ml). In contrast, SPP1 showed a delayed plasma increase (days 8 and 15), and MUC1 and SP-A changes were restricted to BALF. Changes in MMP12 were not detected in BALF or plasma study samples using the available ELISA method, nor were they detected in BALF proteome samples throughout the study.

Together these data suggest that several candidate biomarkers, such as RETNLA, SP-D, LCN2, and MMP7, exhibit significant and consistent changes in plasma and BALF following AZ-1 treatment, underscoring their potential for monitoring lung toxicity.

### Cross-species confirmation reveals consistent lung toxicity mechanisms and supports biomarker translational relevance

To evaluate whether AZ-1-derived biomarker candidates reflect compound-specific or general mechanisms of lung toxicity, we assessed their expression in seven independent datasets. These included: (i) lung transcriptome and BALF proteome datasets from rats treated with a second inhaled compound, AZ-2, known to induce immune cell infiltration and macrophage vacuolation, and (ii) mouse BALF proteome datasets from six established lung injury models, each representing distinct pathological mechanisms relevant for drug-induced toxicity. These models included: bleomycin, causing oxidative stress and pulmonary fibrosis through ROS generation [31, 32], lipopolysaccharide (LPS), activating TLR4 signaling to trigger acute lung injury through pro-inflammatory cytokine release [33], IL33, promoting type 2 immune responses and eosinophilic inflammation [34], *Alternaria* spp., enhancing neutrophil recruitment and Th2-mediated allergic lung inflammation [35], cigarette smoke (CS), driving inflammation through sustained immune activation [36, 37], and elastase, degrading elastin in extracellular matrix, causing lung damage, emphysema-like changes, and inflammation [38]. These datasets allowed assessment of the relevance and generalizability of the AZ-1-derived biomarker panel across species and lung injury mechanisms.

In AZ-2-treated rats, BALF analysis revealed elevated neutrophils, eosinophils, and lymphocytes in the high-dose animals (15mg/kg inhaled dose) (**Fig. 5a**), while the low-dose (4 mg/kg inhaled dose) group showed no increase. Bulk lung transcriptomics identified approximately 30% overlap in differentially expressed genes between high-dose AZ-2 and AZ-1, including *Mmp7, Cd177, Lcn2, Chi311, Cxcl6, Bpifb1, Atp6vod2, Retnla, Cxcl6 and Ccl20*.

**Figure 5.**
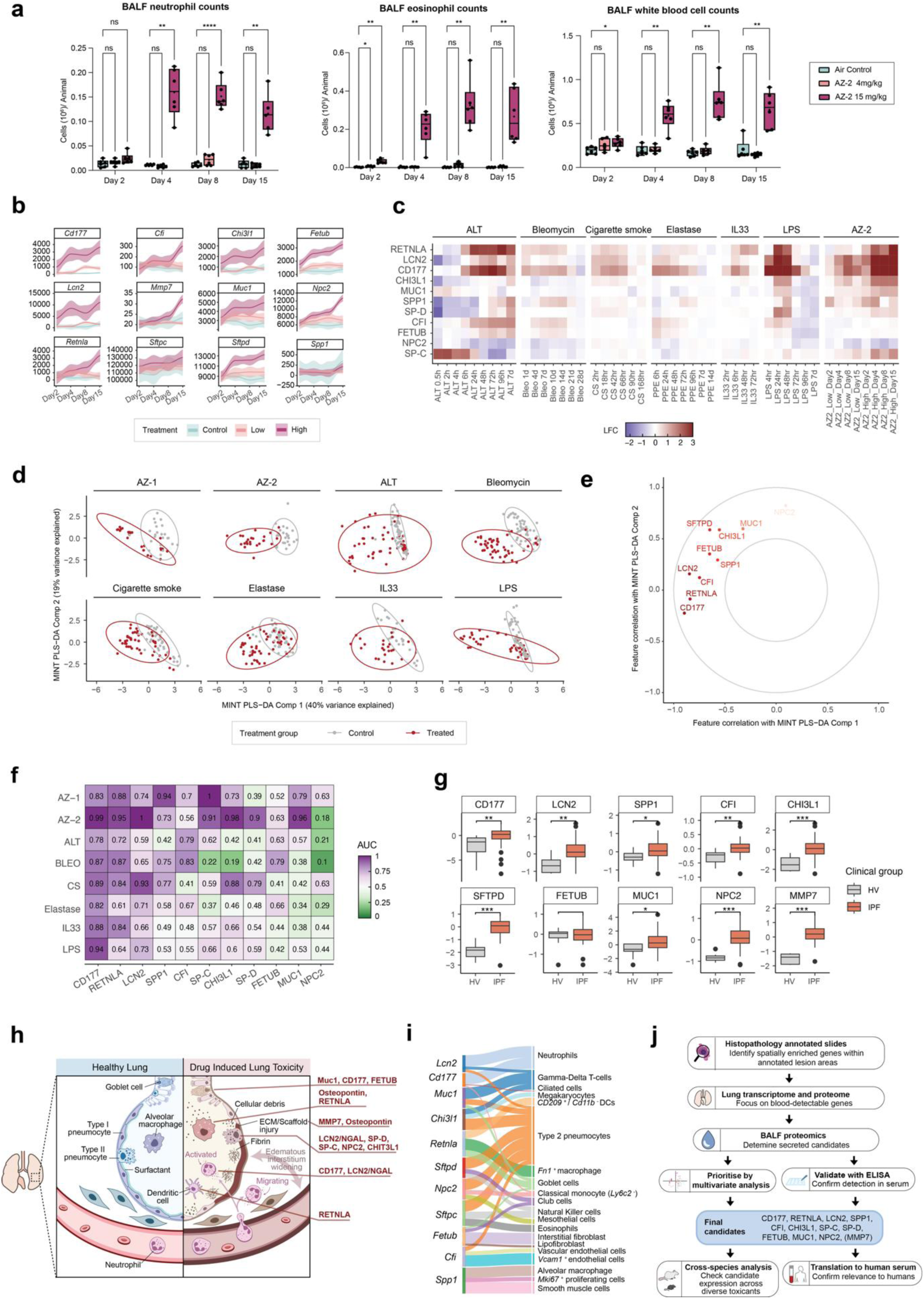
Confirmation of candidates in independent datasets and detectability in plasma samples. **a**. Dose-dependent changes in neutrophils, eosinophils, and white blood cells in BALF after treatment with AZ-2. Significance was tested by two-way ANOVA with Dunnett’s post hoc analysis. **b.** Expression levels of genes corresponding to the 12 AZ-1 derived candidate protein biomarkers in AZ-2 high-dose, low-dose, and control treated lung tissues. **c**. Detection of biomarker candidates in rat BALF treated with AZ-2 or in six independent datasets of toxicant-treated BALF proteomes: LPS, Alternaria, cigarette smoke, IL33, elastase, and bleomycin in mice. **d.** Integration of eight independent datasets (rat BALF treated with AZ-1 and AZ-2, and mouse BALF treated with six lung toxicants) using MINT PLS-DA. Distribution of samples across the first two components is shown and a 95% confidence intervals per treatment group is overlaid. **e.** Correlation of biomarkers with first (x-axis) and second (y-axis) PLSDA components with guidelines of radius 0.5 and 1. **f**. Discriminatory power of candidate biomarkers in distinguishing between toxicity profiles of control and treatment groups in rat BALF treated with AZ-1 or AZ-2, and mouse BALF treated with the six lung toxicants. Color scale indicates area under the receiver operating characteristic (ROC) curve (AUC) for each biomarker in each dataset. Features are ordered by average AUC across all datasets. **g**. Serum protein levels in human samples from IPF patients (n=60) and healthy volunteers (n=9). Protein levels were compared using Wilcoxon rank sum tests with Benjamini-Hochberg multiple testing correction (adjusted p-value < 0.001 = ***, < 0.01 = ** and < 0.05 = *). **h.** Schematic showing cellular sources of the identified biomarkers, with proposed mechanisms and associated toxicity processes **i**. Sankey diagram associating biomarkers with top three cell sources using mouse single-cell atlas data. **j.** Workflow of our comprehensive multi-omics framework for identification of minimally invasive biomarkers through the integrating of spatial and temporal gene expression profiling with proteomic and lipidomic analysis of biofluids.

These overlapping markers primarily play roles in immune cell trafficking, neutrophil degranulation, and airway inflammation [39–41], consistent with the expected in vivo outcome from this compound. Importantly, the majority of the AZ-1-derived biomarker candidates (**Fig. 4d**) were also elevated in the AZ-2 high-dose samples compared to both low-dose and air control groups (**Fig. 5b**), further emphasizing the versatility of the proposed biomarker panel.

To further explore wider applicability and species translation, we examined biomarker expression across six different mouse lung toxicant models. Despite variations in stimuli and timepoints, key proteins such as RETNLA, LCN2, and CD177 were consistently upregulated at early points (24 hours post-stimulation) (**Fig. 5c**). The most significant upregulation for these proteins was observed in response to LPS, *Alternaria* spp., and AZ-2, supporting their value as general indicators of lung toxicity. Some proteins exhibited stimulus-specific associations: CFI was enriched in mice exposed to *Alternaria* spp., bleomycin, and elastase, while SPP1 was elevated in bleomycin-, AZ-2-, and CS-treated samples. This selective response likely reflects each protein’s functional role, such as SPP1’s involvement in fibrosis and tissue remodeling which are key mechanisms in the bleomycin and CS models [42–44].

To explore the spatial expression of the AZ-1-derived biomarkers in an independent injury model, we examined a previously published spatial transcriptomics dataset from bleomycin-injured mouse lungs [45]. Most of our candidate markers were upregulated in bleomycin-treated samples, with elevated expression in fibrotic and/or inflamed areas of the lung. These spatially resolved patterns are consistent with their proposed roles in epithelial remodeling and inflammatory responses and reinforce the relevance of these biomarkers in histologically characterized injury.

To evaluate the consistent changes more systemically, we integrated all eight datasets (AZ-1, AZ-2 and mouse toxicant models) using supervised multi-variate integration (MINT) PLS-DA [46] to find axes of variation best separating control and treated samples across datasets. The first PLS-DA component (**Fig. 5d**) separated treatment groups across all datasets, with the strongest class separation (95% confidence intervals) seen in AZ-1, AZ-2 and bleomycin. We further probed the contribution of individual proteins to this separation by correlating each protein with components 1 and 2 (**Fig. 5e**), which identified CD177, RETNLA, LCN2 and CFI as the strongest contributors.

To further evaluate discriminative power, we performed receiver operating characteristic (ROC) analysis using the BALF proteome datasets (**Fig. 5f**). Strong positive discriminative power (AUC > 0.7) was seen for CD177 across all datasets, and in at least two datasets beyond AZ-1 for seven of the candidates (CD177, RETNLA, LCN2, SPP1, CFI, CHI3L1, SP-D). Some candidates showed model-specificity (e.g. CHI3L1 and SP-C had AUC < 0.5 in the bleomycin model), indicating potential for toxicity-type-specific biomarker applications. In contrast, CD177, RETNLA and LCN2 demonstrated strong (AUC > 0.7) and consistent performance across most models, positioning them as robust generalizable biomarkers.

We next assessed the value of combining candidates into panels. In each dataset, optimal thresholds were calculated for each protein to separate between control and treated samples, and a majority voting approach was applied across several feature sets to classify samples in an unseen test set. Individual features were used, as well as incremental feature sets taking the top 2, 3, 4, … *n* features according to average individual AUC across all datasets (**Fig. 5f**). Under this assessment, CD177 remained the best feature by test accuracy, followed by CD177+RETNLA+LCN2 (Top 3) and CD177+RETNLA (Top 2) then the top n=7, 4, 5 and 9 features which all perform better than any individual features apart from CD177. This highlights the potential power of these biomarkers in combination as a panel for clinical application. MMP7 was not present in the dataset for analysis.

### Mechanistic biomarkers demonstrate human translatability

Given the mechanistic overlap between drug-induced lung toxicity and chronic interstitial lung disease, we next assessed the presence of AZ-1-derived biomarker candidates in human samples from patients with idiopathic pulmonary fibrosis. IPF is partially characterized by immune activation and release of pro-inflammatory and pro-fibrotic cytokines, mirroring features of drug-induced lung injury. This pathophysiological similarity suggests that serum derived from IPF patients may serve as a valuable resource for evaluating the human translational potential of pre-clinical biomarker candidates.

We analyzed serum samples from 9 healthy volunteers (HV) and 60 patients diagnosed with IPF. Of the 12 AZ-1-associated biomarkers identified in preclinical models, 10 were detectable in the human dataset (**Fig. 5g**). All but fetuin B (FETUB) were significantly upregulated in IPF patients relative to healthy controls. These findings support the relevance of the identified markers to human lung injury and strengthens their potential utility in monitoring drug-induced lung injury in clinical settings. To further investigate their spatial relevance in a chronic disease setting, we examined a previously published spatial transcriptomics dataset from IPF patient lungs [45]. In this dataset, *SPP1*, *CHI3L1*, *MMP7* and *CFI* were upregulated in IPF samples compared to healthy controls, with *SPP1*, *CHI3L1*, and *MMP7* particularly enriched in regions with extensive fibrosis.

Taken together, our studies identified 28 transcriptomic biomarker candidates in AZ-1-treated lung tissue, with 18 confirmed at the protein level in BALF in our study samples. Of these, 12 proteins demonstrated high discriminative power for identifying toxicity in BALF proteomics data (**Fig. 5h-j**). Five markers—LCN2, RETNLA, SPP1, SP-D, and MMP7—were also detectable and predictive in plasma, offering potential for less invasive monitoring. Seven biomarkers demonstrated robust discriminative power (AUC > 0.7) across seven additional preclinical lung injury models. Panel-based prediction consistently outperformed individual markers, highlighting the benefit of using these biomarkers in combination. Furthermore, we confirmed their presence and detectability in human plasma and their elevation under disease conditions where lung toxicity is evident. Given this observation, it is worth highlighting that assessing drug-induced lung injury using these biomarkers in healthy volunteers was necessary, as disease-related effects might alter biomarker levels and influence the evaluation of drug-induced effects. Altogether, we identified a panel of 12 protein biomarkers with translational potential for diagnosing and monitoring lung injury preclinically as well as in humans.

## Discussion

Our study addresses the critical unmet need for early, non-invasive biomarkers to improve safety assessment in inhaled drug development. By integrating spatial and temporal gene expression from rat lung tissue with proteomic profiling of biofluids, we established a comprehensive multi-omics framework to uncover molecular signatures of drug-induced (lung) injury.

This approach enabled the identification of a panel of 12 biomarker candidates with spatial, temporal and mechanistic relevance to drug-induced lung injury, particularly inflammation and epithelial remodeling. The strength of our framework lies in its stepwise design. Starting with spatial transcriptomics to identify tissue region- and cell type-specific molecular changes, followed by targeted confirmation of these signals in BALF and ultimately in circulation (**Fig. 5h-j**). Our results also indicate that utilizing a multiplexed biomarker panel approach enhances predictive performance, capturing the diverse mechanisms of lung toxicity more effectively than single biomarkers. Furthermore, confirmation in human serum from patients with IPF confirmed the presence of several candidate biomarkers, supporting their detectability in humans and their relevance to lung injury.

Many of the identified candidates have established roles in lung injury and inflammation and are clinically recognized markers of tissue damage. For example, LCN2 (NGAL) is a rapid responder to tissue injury and correlates with inflammation and fibrosis severity in chronic lung diseases such as COPD and IPF [47]. CD177, a neutrophil glycoprotein, is upregulated in inflammatory lung conditions, including ARDS, with anti-CD177 treatment shown to reduce lung inflammation in animal models [48]. Surfactant dysfunction, indicated by altered phospholipid composition, is a hallmark of lung injury, with elevated SP-D levels correlating with epithelial damage and disease severity[49, 50]. MMP7 plays a pivotal role in extracellular matrix remodeling and is consistently elevated in fibrotic lung diseases [51, 52]. Osteopontin (SPP1) is increased in lung infections and viral injury, with circulating levels correlating with severity in COVID-19 patients [53, 54]. Moreover, SPP1+ macrophages contribute to fibrosis in both humans and mice [55]. Collectively, these findings reinforce the biological relevance and translational potential of the biomarker candidates identified here, for drug-induced lung toxicity monitoring.

Our study provides compelling evidence for mechanistic understanding and translational biomarkers; however, several areas present opportunities for further development. Our initial single-compound approach was strengthened by subsequent validation across multiple toxicants, confirming broad applicability. Species-specific assay availability limited full validation in plasma for all candidates. RETNLA, while lacking a human ortholog, remains valuable for preclinical assessment. Future studies should expand compound diversity and refine biomarker specificity.

Despite these limitations, our spatial multi-omics framework advances understanding of lung toxicity mechanisms and identifies early biomarkers with spatial and circulatory relevance. By linking cell type-specific responses to secreted biomarkers, our study offers insights to guide the development of predictive in vitro models and non-animal screening methods. This approach holds the promise of enhancing preclinical safety assessment and accelerating respiratory drug development, with potential regulatory impact in biomarker qualification and clinical monitoring.

## Methods

### In Vivo studies

Animal experiments associated with AZ-1 and AZ-2 were conducted in an AAALAC-accredited AstraZeneca facility in Gothenburg, Sweden, with ethical approval (no. 2020-002853). Male Wistar Han rats (10 weeks old, ∼250 g), group housed under standard conditions, were acclimatized and trained for inhalation exposure. Rats received daily inhaled doses of AZ-1 (2 and 15 mg/kg) or AZ-2 (4 and 15 mg/kg) for up to 14 days; control groups were exposed to air. AZ-1 was administered via a Wright Dust Feed (WDF) and AZ-2 with a Rotating Brush Generator (RBG), both producing dry powder aerosols delivered to the inhalation chamber. All doses are presented as total inhaled dose; the actual lung deposited dose is 10% of the inhaled dose. Aerosol concentration and particle size were verified in vitro prior to each exposure by measuring drug deposition on a glass fiber filter at the inhalation point. Additional groups received single high doses for spatial transcriptomics and metabolomics analyses.

Studies with additional six lung toxicants were performed in mice. Female C57BL/6 mice (8-12 weeks, 18-22g, Charles River Laboratories) were housed in AAALAC-accredited facilities with standard conditions (12h light-dark cycle, 21±1°C, 55±15% humidity) and environmental enrichment. Animals were monitored daily with defined humane endpoints. Experiments were approved by Gothenburg Ethics Committee (BLM, CS, IL33, LPS) or UK Home Office (ALT, ELAST), conforming to EU Directive 2010/63/EU and institutional standards.

After 5-day acclimatization, mice were challenged with various lung toxicants under isoflurane anesthesia (except cigarette smoke): BLM - 40µg bleomycin (Apoleomycillo Scientific, BI3543, Chemtronica Sweden) oropharyngeally (30µl); ALT - 25µg Alternaria alternata (Greer, XPM1D3A25) intranasally (50µl); ELAST - 10U/kg porcine pancreatic elastase (Sigma, E1250) intratracheally (80µl); CS - cigarette smoke (Kentucky Research cigarettes, 1R6F) exposure (60min, twice daily); IL33 - recombinant mouse IL-33 (Peprotech, 210-33) intranasally (50µl); LPS - lipopolysaccharide (Sigma, L2630, serotype 0111:B4) aerosol (30min). Controls received appropriate vehicles. At specified timepoints, animals were terminally anesthetized with sodium pentobarbital. BALF was obtained and subsequently centrifuged and supernatants aliquoted and stored at −80°C.

### Tissue and biofluid sample collection

Samples were collected at four time points (days 2, 4, 8, and 15), 24 hours post-final dose to capture both early and late toxicological effects.

### Clinical chemistry samples

A minimal volume of 500 μL of blood was collected from vena cava into Microvette 500 LiHep tubes (Sarstedt). Tubes were gently inverted 10 times and placed on ice. Plasma was prepared within 30 minutes of sampling by centrifugation (1500×g, 10 min, 4°C). The samples were stored at −80°C and shipped to Charles River Laboratories, Edinburgh, UK, for analysis of plasma chemistry readouts.

### Broncho-alveolar lavage and preparation of BALF

Broncho-alveolar lavage was performed by manual perfusion of the whole lung. A volume of 4 mL room temperature PBS was injected into the lung via the exposed trachea, and the fluid was then recollected by slow aspiration into the syringe. This procedure was performed twice.

### Plasma isolation

A volume of 6-8 mL of blood was collected from vena cava into S-Monovette Li-Hep tubes (Sarstedt). Plasma was prepared within 30 minutes of sampling by centrifugation (1500×g, 10 min, 4°C) and stored at −80°C.

### Tissue samples for histology

Left lung lobes were excised and fixated in 4% formaldehyde solution for 48 hours. Fixed tissues were stored at room temperature until further processing.

### Tissue samples for spatial transcriptomics and MSI

A volume of 0.4 mL heparin was administered intracardially through the diaphragm. The abdominal aorta was cut to facilitate exsanguination, followed by an incision in the apex of the heart. The pulmonary circulation was perfused with 37°C saline, followed by 37°C low melting agarose solution. The lung was then inflated through the trachea with 2 mL of agarose solution and tied off. The tissue was excised, snap-frozen in pre-chilled isopentane/dry ice slurry for 30 seconds, and stored at −80°C.

### Tissue samples for RNA and protein analysis

For RNA analysis, three pieces of the right inferior lung lobe was cut off (approx. 10 mg each, total max 30 mg), placed in DNA LoBind tubes (Eppendorf) pre-filled with 1 mL RNAlater (Invitrogen), and stored at 4°C overnight. The samples were then transferred to a new tube and stored at −80°C until RNA extraction.

For protein analysis, a minimum of 5 mg of the right inferior lung lobe was excised, rinsed with PBS to remove any blood, and snap-frozen in liquid nitrogen. Samples were stored at - 80°C until analysis.

### Histological processing and assessment

Formalin fixed lung tissues were prepared in a tissue processor (MAGNUS, Milestone Medical, Sorisole, Italy). Samples were dehydrated in increasing concentrations of ethanol, then placed in xylene substitute, infiltrated with liquid paraffin, and embedded into paraffin blocks. Blocks were cut into ultrathin 4 µm tissue sections using a microtome (RM2165 Leica Microsystems, Triolab AB, Gothenburg Sweden) and placed on glass slides (Superfrost Plus, Thermo Scientific, Braunschweig Germany). Tissue sections were then stained with hematoxylin and eosin in a Leica ST5020 multistainer (Leica biosystems, Triolab AB). Stained slides were scanned at 40x magnification using a brightfield scanner (Aperio AT2 Leica scanner, Leica Biosystems, Braunschweig Germany).

Histopathological assessments were performed on H&E-stained tissue sections using the Loupe Browser (10x Genomics) software and were manually annotated based on tissue morphology. The human lung data were classified into ‘blood vessel’, ‘large airway’, and ‘suspect inflammation’, where ‘suspect inflammation’ was distinguished as areas with dense aggregations of immune cells

### Spatial Transcriptomics

Fresh-frozen agarose-inflated rat lung tissues were cryosectioned at a thickness of 10-12 µm, with the cryostat temperature set to −20°C and the knife to −10°C. Several tissue sections were collected and stored at −80°C for RNA extraction and quality assessment. RNA integrity was assessed using a 5300 Fragment Analyzer (Agilent) and RIN values were >9 for all samples. Consecutive lung tissue sections were then immediately positioned onto a Visium Gene Expression slide (for spatial transcriptomics) and a Superfrost slide (for MSI), which was stored at −80°C for further processing.

#### Sample preparation

Tissue fixation, staining, and imaging were conducted following the Methanol Fixation, H&E Staining, and Imaging Visium protocol provided by 10X Genomics. Stained sections were imaged at 40X magnification using an Aperio Digital Pathology Slide Scanner (Leica Biosystems), ensuring visual data for the spatial analyses. Library preparation for sequencing was performed in accordance with the Visium Spatial Gene Expression User Guide (Rev E; 10X Genomics). Tissue sections were permeabilized for 15 minutes, based on prior optimization using the Visium Tissue Optimization kit, followed by cDNA amplification and indexing.

The prepared libraries were pooled and sequenced with a 1% PhiX spike-in on a NovaSeq 6000 (Illumina) platform using an S4 flow cell. Loading concentration was 300pM. The sequencing yielded between 177-299 M reads per sample. An average of 4,179 spots per sample were covered by tissue, with an average of 57,000 reads per spot across samples.

#### Spatial Lipidomics by Desorption electrospray/ionization mass spectrometry imaging (DESI MSI)

The Fresh-frozen agarose-inflated rat lung tissues thaw mounted onto a superfrost slide were used for the MSI based spatial lipidomics experiment. DESI MSI was carried as recently described in Sushentsev et al.[56] Analyzes were performed at spatial resolutions of 60 μm in negative ion mode and mass spectra were collected in the mass range of 80−900 Da with mass resolving power set to 70000 at m/z 200. Haematoxylin and eosin (H&E) staining was performed post-analysis on the same tissue sections and the stained sections were imaged at 20x with Aperio CS2 digital pathology scanner (Aperio Tech., Oxford, UK). The .imzml data files were imported into SCiLS Lab software (Bruker Daltonics, Germany, 2022a MVS). Bisecting k-means segmentation was performed within SCiLS Lab to create two regions of interest (ROIs): tissue and background (area outside the tissue). Individual samples were then segmented using the tissue/background ROIs with some manual alterations. Individual whole tissue-level Root-Mean Square (RMS) normalised peak intensities were extracted using the SCiLS Lab API, which yields a pixel-by-peak table with associated metadata. This table was later used as input to the MAGPIE pipeline [Williams 2025, https://www.biorxiv.org/content/10.1101/2025.02.26.640381v1.full] to co register with Visium spatial transcriptomics data. Manual landmarks were selected through the MAGPIE Python Shiny app which were then used to first coregister MSI data coordinates to the H&E image from the MSI section then to the Visium H&E image. MSI pixels were then aggregated to Visium spots and compared to pathology annotations. The peaks were annotated against compounds in the KEGG database using [M-H]- and [M+Cl]- adducts with a mass tolerance of 5 ppm.

### Mass spectrometry analysis

Total lipid extraction from both plasma and BALF samples was performed using a modified version of the method from Sarfian et al.(56). Specifically, samples were diluted in analytical grade water at a ratio of 1:3 prior to overnight protein precipitation by adding 3× volumes of 100 % isopropanol solution containing equiSPLASH (330731, Avanti, 1:50 final) and CUDA (10007923, Cayman Chemical, 3 µM final) internal standards. After centrifugation, the supernatant was retained and was mixed with 100% acetonitrile solution at 2:1 ratio (final ratio 3:2:1 IPA:ACN:H2O) and centrifuged again to ensure no particulates were carried over to the mass spectrometry vial. One µL of supernatant was injected for liquid chromatography-mass spectrometry (LC-MS) analysis.

Lipidomics analysis was performed using a Vanquish UHPLC coupled to an Orbitrap IQ-X mass spectrometer (ThermoFisher). Chromatographic separation was performed on an Acquity CSH C18 column (1.7μm, 2.1mm × 100mm, Waters Corporation). Mobile phases A and B were 60:40 acetonitrile:water with 10mM ammonium formate and 0.1% formic acid and 90:10 isopropanol:acetonitrile with 10mM ammonium formate and 0.1% formic acid, respectively. Flow rate was maintained at 0.6 mL/min for the entire duration of the LC method. The gradient was increased from 15% B to 30% B over 2 min, 30% B to 48% B over 0.5 min, 48% B to 82% B over 8.5 min, 82% B to 99% B over 0.5 min, held at 99% B for 0.5 min, and equilibrated at initial conditions for 3 minutes. The column compartment was maintained at 65°C. Lipid detection was performed by MS1 polarity switching analysis of each individual sample with an Orbitrap resolution of 30,000 from 150-1200 m/z. Lipid identification was performed by iterative MS/MS using stepped HCD fragmentation and the AcquireX DeepScan workflow (ThermoFisher) on a representative pooled sample from each specimen and treatment. Lipidomics data was analyzed using MS-DIAL v 4.9. Peak detection, adduct assignment, identification, and alignment were performed in MS-DIAL. Lipid annotations were performed using the *in-silico* fragmentation spectral library provided with MS-DIAL. The features aligned across samples and treatments, generated by MS-DIAL, were subjected to normalization by total ion chromatogram. Lipids were filtered based on signal intensity (>3X blank) and RSD >30%. Further filtering removed lipids classes that were < 5% of the total lipid intensity.

### Tissue transcriptomics

#### Total RNA extraction

Total RNA was purified from lung tissue samples with RNAdvance Tissue^TM^ kit by Beckman Coulter Life Sciences (Catalog number A47943) in a 96-well plate on a Beckman Biomek i7 liquid handler instrument. The quantity and quality of RNA samples was then assessed on a Fragment Analyzer 5300 (Agilent Technologies). The data obtained was analysed using ProSize data analysis software to derive the concentration and RNA quality numbers (RQN) for individual samples. All samples were deemed of sufficient quality and quantity for library construction.

#### RNA-Seq library construction and sequencing

The double-stranded cDNA obtained with Poly (A)-mRNA enrichment (500 ng total RNA input) was subjected to library preparation using an mRNA HyperPrep Kit (Kapa Biosystems, MA, USA), and prepared according to the manufacturer’s protocol. Preparation was performed automatically on a Tecan Fluent. The quality of the libraries was determined using a 5300 Fragment analyzer with the standard sensitivity NGS kit (Agilent). Libraries were then pooled and denatured according to Illumina’s recommendations and sequenced on a NovaSeq6000 on a PE50 configuration, to achieve a mean coverage of ∼20M reads per library.

#### RNA-seq data processing and analysis

The initial quality control (QC) was performed using FastQC and MultiQC [57], which indicated some adapter contamination and variable sequencing depth; as a result, all reads were trimmed to 45nt and subsampled to 30M reads, then aligned to the *R. norvegicus* genome (MRatBN7) using STAR v2.7.10b [58] and quantified using featureCounts (Rsubread v2.8.2) [59]. The resulting expression matrices for each batch were denoised using noisyR [60] using a signal/noise threshold of 20 and quantile normalised [61] further QC and analyses were performed using bulkAnalyseR v1.1.0 [62]. Differential Expression analysis with edgeR [63] used |log2(FC)| > 0.5 and 1.5 and a Benjamini-Hochberg-adjusted p-value of < 0.05. Resulting genes were annotated using gene set over-representation analysis through gprofiler2 [64] and Ingenuity Pathway Analysis [16].

### Tissue proteomics

Lung tissue samples were pulverized using a Covaris CP02 cryoPREP Automated Dry Pulverizer and lysed in PreOmics iST buffer. Protein quantification was performed using the BCA Protein Assay Kit, followed by denaturation at 95°C for 10 minutes. Proteins were digested overnight with Lys-C and trypsin in a 1:50 enzyme-to-protein ratio. The resulting peptides were desalted according to the PreOmics iST protocol and vacuum-centrifuged to dryness, then resuspended in a solution of 2% acetonitrile and 0.1% formic acid. Peptides were quantified using the Quantitative Colorimetric Peptide Assay before being fractionated into 24 fractions via high pH reversed-phase chromatography on an ACQUITY UPLC Peptide BEH C18 column. The LC-MS/MS analysis was conducted on a Thermo UltiMate 3000 system coupled to either an Orbitrap Eclipse Tribrid or Orbitrap Exploris 480 mass spectrometer. The mobile phases consisted of water with 0.1% formic acid (Phase A) and 90% acetonitrile with 0.1% formic acid (Phase B). Peptides were first trapped on an Acclaim PepMap 100 C18 trap column before separation on an Aurora Ultimate column using a nonlinear gradient. Data acquisition involved multiple MS1 and MS2 scans using high field asymmetric waveform ion mobility spectrometry (FAIMS) for enhanced peptide separation. The mass spectrometer operated in positive polarity, with specific settings for spray voltage, capillary temperature, and RF levels. Dynamic exclusion was set for precursor ions to optimize detection. Project-specific libraries were generated from the acquired data, which were analyzed using Spectronaut software, maintaining a 1% false discovery rate for both precursor and protein identification. Static modifications included carbamidomethylation of cysteines, while dynamic modifications accounted for N-terminal acetylation, oxidation of methionine, and deamidation of asparagine and glutamine.

### Biofluid proteomics

10uL of BALF was used for analysis. BALF samples were processed using the EasyPep kit (Thermo Scientific), following the manufacture’s protocol, dried using rotary evaporation, and stored at −80°C. Plasma was depleted using TOP14 resin (Thermo Scientific), dried using rotary evaporation, and stored at −80°C prior to processing using the EasyPep Kit. Following EasyPep, dried peptides were resuspended in 0.1% formic acid for BALF and plasma to achieve a nanodrop OD <1. 250ng of peptide (in 60µL) was loaded onto Evotips for mass spectrometry analysis. Peptides were separated using an EvoSep followed by detection on a Bruker timsTOF ProII with data acquired in DIA mode. BALF and plasma samples with high blood contamination (±2SD on the hemolysis index) were removed from downstream analysis [65] Data analysis was performed using a directDIA approach in Spectronaut (Biognosys) with the rat Uniprot database FASTA file from September 2022 and the following settings: imputation strategy – none; cross-run normalization – true; normalization filter – none; normalization strategy – automatic; and row selection – automatic.

Normalized protein abundances exported from Spectronaut were first assessed for batch effects and outliers by principal-component analysis and unsupervised hierarchical clustering. Proteins quantified in > 50 % of samples were retained; remaining missing values were imputed with the pcaMethods package (svdImpute function) in R.

#### Computational methods for circulatory proteomics

After normalisation and imputation, one air control sample (BAL.75) was removed as an outlier based on a 99% confidence interval on all air control samples. Principal component analysis was performed using potential candidates from Visium and bulk transcriptomics on scaled and centred BALF proteomics data using prcomp function and visualised using factoextra. The contribution of individual proteins to PC1 and PC2 were further analysed to select discriminative biomarkers. The same problem was approached from a different angle, using PLS-DA as implemented in mixOmics [66] on scaled data with 2 components separating based on the treatment group (air control, low dose or high dose). The contribution of individual proteins was analysed based on correlation with the PLS-DA components (using the plotVar function).

#### Proximity extension Assay

Serum protein concentrations were measured using the OLINK HT Target (OLINK Proteomics, Uppsala, Sweden) following the manufacturer’s instructions. Plasma samples from 60 idiopathic pulmonary fibrosis (IPF) patients, and 9 healthy controls were analyzed using the OLINK HT panel (∼5,440 proteins). All patients and healthy volunteers signed informed consent prior to inclusion to the study. All experiments with human tissue samples were performed under protocols approved by the Institutional Review Boards at Hannover Medical School (IRB #2923-2015). Briefly, 2 µL of serum was incubated with paired oligonucleotide-labeled antibodies, enabling proximity-dependent DNA hybridization and polymerization (Proximity Extension Assay, PEA). Extension products were pre-amplified by PCR and quantified using NGS on a NovaSeq 6000 platform (Illumina). Data were normalized and quality controlled with OLINK NPX Manager software, and protein levels were reported as NPX units (log2 scale). Samples and controls were randomized, and all procedures followed OLINK protocols. Group comparisons were performed using two-sided Wilcoxon tests.

#### Enzyme-Linked Immunosorbent Assays (ELISA)

ELISA kits for SP-D (NBP2-76687), SP-A (NBP2-76694), MMP7 (NBP3-06896), MMP12 (NBP3-06764), and Muc1 (NBP2-74991) were obtained from Novus Biologicals, while the rat OPN ELISA kit (ERA46RB) was purchased from Thermo Fisher Scientific. Assays were performed in accordance with manufacturers’ instructions. Briefly, antibody-precoated plates were washed and blocked before addition of standards and appropriately diluted samples, each loaded with three biological replicates. After incubation at room temperature, detection antibodies were added, followed by the addition of a colorimetric substrate. The enzymatic reaction was stopped, and absorbance was measured at 450 nm. Protein concentrations were determined by comparing sample absorbance values to the standard curve. For statistical analysis, analyte intensities were compared across dose levels at each timepoint using the Kruskal-Wallis test, followed by Dunn’s post-hoc test and Benjamini-Hochberg correction for multiple testing.

#### Meso Scale Discovery (MSD) assays

Rat serum samples were analyzed for LCN2 (K1535ER) and RETNLA (K152ZOR) using Meso Scale Discovery (MSD) assays. Samples were diluted 1:100, and standards were prepared according to the manufacturer’s instructions. Standards and diluted samples were added in triplicate to prewashed plates and incubated at room temperature with gentle shaking (750 rpm) for 1–2 hours, followed by washing. Detection antibody was then added, and plates were incubated for another hour with shaking, followed by a final wash. MSD read buffer was applied, and electrochemiluminescence signals were recorded using an MSD Sector Imager. Analyte concentrations were calculated by comparing sample signals to the standard curve in Methodological Mind software. Intensities were compared between dose levels at each timepoint using the Kruskal-Wallis test, with post-hoc Dunn’s test and Benjamini-Hochberg correction for multiple testing.

#### Candidate prioritisation in circulatory proteomics

To assess the generalisability of signals across multiple lung toxicants, we integrated eight datasets (rat BALF treated with AZ-1 or AZ-2 (air control vs high dose), and mouse BALF treated with LPS, Alternaria, cigarette smoke, IL33, elastase, or bleomycin) using multivariate (MINT) PLS-DA from mixOmics [46, 66] with 2 components to separate between control and treated samples. The contribution of individual proteins was analysed based on correlation with the PLS-DA components (using the plotVar function).

The discriminative power of each candidate individually was further assessed using ROC analysis (using pROCpackage) [67] comparing control and treatment groups in the same eight datasets across all timepoints, specifying that the treatment expression should be higher than the control expression [67]. The area under the curve (AUC) was reported for each dataset and each protein.

To further assess the combined power of candidates selected from BALF proteomic analysis, combinations of features were tested using a multivariate thresholding approach. Each dataset was split into a training (75%) and testing (25%) set, ensuring even representation of treatment groups. Using the training set, optimal thresholds were calculated using Youden’s J statistic with the coords function from pROC [67]. These thresholds were applied to both training and test datasets and combined classifications for each sample were established were based on whether > 50% of features/proteins (among several tested feature sets) passed the threshold. Training and test accuracies were calculated by comparing these combined classifications against true treatment group labels. The tested feature sets were (i) individual proteins, (ii) the top n (n=2,3,4,) features according to AUCs from ROC analysis described previously.

### Statistical analyses

Statistical methods for each analysis are detailed in their respective method sections.

## Reporting summary

Further information on research design is available in the Nature Portfolio Reporting Summary linked to this article.

## Data availability

All Visium datasets, including raw fastq files, SpaceRanger output, and high-resolution images, needed to replicate and further build upon the presented analyses will be deposited to GEO upon publication. The raw and processed bulk tissue RNA-sequencing data for the samples will also be deposited to GEO.

## Acknowledgements

This work was supported by the Swedish Foundation for Strategic Research (grant no. ID18-0094 supporting L.F.), the MRC DTP iCASE studentship programme (G117817 supporting E.C.W.), and AstraZeneca. IM was funded by the Wellcome Trust [203151/Z/16/Z] and the UKRI Medical Research Council [MC_PC_17230]. We would like to thank the animal science team for their invaluable support with the in vivo studies and termination, and Linnea Johansson and Kelley Patten for their assistance with the termination procedures. Our gratitude also extends to the team who provided the idiopathic pulmonary fibrosis (IPF) dataset. We also extend our gratitude to the following people who were instrumental in generating samples that were essential for the success of this study: Alison Dodd, Thomas Volckaert, Theresa Andreasson, Marie Jönsson, Gopinath Kasetty, and Helena Douglasson. We also thank Lisa Öberg for bioinformatics support across all studies relevant to the mouse BALF dataset.

## Author contributions statement

M.M.M., A.O. and J.J.H. and P.Å conceived the study and secured resources. M.M.M co-ordinated and led the project with input from A.O., P.Å., and K.O. M.M.M., C.D.P., E.L.A., J.C., A.A.I., M.O.L., and J.L.T. designed the experiments. S.O. and R.H.A. planned and conducted the animal dosing studies and performed the inhalation exposures. M.M.M. and M.O.L. collected the lung and biofluid samples. E.S., A.C supported the processing of histopathological samples and L.S. conducted the histopathological evaluations and annotations. M.O.L. processed and sectioned tissue samples for spatial transcriptomics and mass spectrometry imaging. M.O.L. performed the spatial transcriptomics experiments. G.H. and S.J performed the spatial lipidomics and C.D.P. performed mass spectrometry. J.L.T. and J.L. generated the bulk RNA-seq. A.A.I. and E.M. generated the proteomics data. B.P.K. pre-processed the spatial transcriptomics data. G.H. analysed the spatial lipidomics data, and A.J., A.A.I., and E.M. analysed the proteomics data. E.C.W., M.M.M., A.J., D.P.C., E.J.A, S.H and J.C. analysed the biofluid data from BALF and plasma. E.C.W. carried out computational analyses of the spatial and bulk transcriptomics, as well as tissue and BALF proteomics data, and performed the integration of spatial and bulk multi-omics with input from J.T. and L.F. E.C.W. performed visualization and created final figures and illustrations. M.M.M., M.O.L., and E.C.W. drafted the manuscript, with critical input from all authors. I.M., J.T. and PS supervised the data analyses. P.F and J.J provided resource and critical input to complete in vitro assay. D.C. and A.B. provided the mouse lung inflammation dataset. A.A. and A.P. generated IPF dataset. All authors reviewed and approved the final manuscript.

